# Lipid droplet shape and tendency towards budding: insight from theory and molecular simulations

**DOI:** 10.64898/2026.07.13.736997

**Authors:** Vincent Nieto, Jackson Crowley, Francois Deslandes, Abdou Rachid Thiam, Lionel Foret, Luca Monticelli

## Abstract

Lipid droplets (LDs) are cellular organelles responsible for lipid storage and metabolism. The mechanism of biogenesis of LDs involves phase separation of neutral lipids from the surrounding phospholipids, which generates oil lenses embedded in lipid bilayers, also known as nascent LDs. As nascent LDs grow, at some point they bud out of the bilayer, forming nearly spherical droplets. Nascent LDs have different propensity to bud, and it has been proposed that their shape provides information on such propensity; however, LD shape is difficult to determine experimentally.

Here we studied the shape of lipid droplets using MD simulations at the coarse-grained level, and compared it to the predictions by an established theory. Our general system setup features an oil lens embedded into a flat, periodic bilayer. We found that the shape of simulated nascent LDs resembles a spherical cap (i.e., it has constant curvature over most of the surface), in excellent agreement with the theory, already for very small droplet sizes. The aspect ratio (height/radius) of nascent LDs increases with increasing LD volume, increasing membrane softness, and increasing surface tension between oil and water, also in agreement with theoretical predictions; however, it remains lower than 1 (i.e., the ratio for a sphere) for LDs of up to 40 nm in diameter. Fitting the simulated LD shapes with a theoretical shape equation suggests that a non-zero surface tension is present in both the monolayer and in the bilayer region. The existence of a relatively high surface tension in the bilayer region is confirmed by local stress calculations, and indicates that the periodic system setup does not reproduce the properties of nascent LDs in the endoplasmic reticulum, where the bilayer tension is two orders of magnitude lower. However, the simulations provide a microscopic view into the properties of droplet embedded vesicles.

## Introduction

Lipid droplets (LDs) are generated from the endoplasmic reticulum (ER) membrane, and perform their functions in lipid storage and metabolism when they are budded out of the ER membrane, towards the cytosolic compartment (1–3) . For this reason, it is very important to understand what determines the direction of budding and the propensity of nascent LDs to bud. Theoretical models can provide useful insight into the driving forces for LD budding (4–6). Available theories are based on a continuous media description (Helfrich membrane theory (7)) and take into account mechanical properties of membranes, such as their elastic energy, the surface tension of the interfaces, and the spontaneous curvature of each membrane leaflet. Thiam and Foret developed a theory describing the mechanics of LD budding starting from a nascent LD, i.e., an oil lens sandwiched between the ER leaflets (4). The theory is based on a continuous media description and takes into account the elastic energy of the membrane, as well as the surface tension of monolayer and bilayer membranes. A compact description of the theory is provided by the following equation:

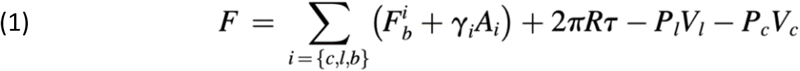

with

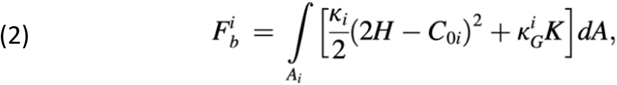

where the first term in equation 1 represents the elastic energy of the three portions of the membrane; *i = [c,l,b]* indicates the cytoplasmic monolayer, the luminal one, and the bilayer respectively; the second term represents the surface tension of the same portions of membrane; *τ* and R are the line tension and radius of the contact line, respectively; and *P*_*c,l*_ and *V*_*c,l*_ are the pressures and volume changes of the cytosol and lumen, respectively. Equation 2 indicates that the elastic energy depends in general on *k*_*i*_, the bending rigidity; *C*_*0i*_, the spontaneous curvature; and *k*^*i*^_*G*_, the Gaussian bending rigidity of the membrane *i* and on the local mean and Gaussian curvatures *H* and *K* (4). The shape equation above must be complemented by appropriate boundary conditions at the three-phase contact line, where the two monolayers and the bilayer meet. These boundary conditions are: (i) geometric continuity of the membrane position and slope; (ii) force balance (analogous to the Neumann triangle condition), which relates the monolayer and bilayer tensions to the contact angle; and (iii) torque balance, which requires continuity of the mean curvature across the contact line. This latter condition ensures that the bending moments exerted by the monolayer and the bilayer are balanced at their junction. Together, these boundary conditions fully specify the shape of the nascent LD for a given volume and set of mechanical parameters. By minimizing the free energy expression in equation 1, considering estimates for the physical constants (surface tensions and elastic constants), the theory allows to compute the shape of an LD embedded in the ER for any volume of the LD. For example, the same paper reports the shapes of nascent LDs for different values of LD volume and surface tension of the monolayer and bilayer regions (Figure 2 in ref. (4)). According to this theory, oil lenses are flat at small oil volume, when the typical droplet size is smaller than the elastic length 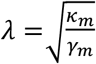 where *κ*_*m*_ is the bending rigidity of the monolayer at the droplet interface and *γ*_*m*_ is surface tension. At larger size, the surface tension dominates and imposes a nearly spherical shape. In this situation, the lens can become unstable leading to the spontaneous formation of a spherical protrusion because of the monolayer deformation near the droplet edge that gives rise to an effective line tension. The analysis of LD shape is particularly interesting in the context of LD biogenesis, as shape is a proxy for the tendency of a nascent LD to bud out of the bilayer membrane: nascent LDs with a shape close to a sphere have a high tendency to bud, while flatter ones tend to remain embedded in the bilayer membrane (8). Such correspondence between shape and tendency to bud has been observed experimentally, both in synthetic model systems (9) and in biological systems (10).

As stated above, theories are based on a continuum description of the system and contains several approximations (4–6). On the other hand, the early stages of LD budding are difficult to study experimentally, and obtaining high-resolution images of budding LDs is very challenging for current microscopy techniques. To provide a microscopic interpretation of the theory and a molecular view into the mechanism of budding, here we performed molecular dynamics simulations at the coarse-grained level for nascent LDs of different size. We compared LD shapes predicted by the theory and calculated directly from large scale MD simulations, and found an excellent match. More specifically, we found that the shape of simulated LDs has constant curvature, i.e., it resembles a spherical cap, already for droplets of very small size (10 nm), indicating that the shape is largely dominated by surface tension, as predicted by the theory. On the other hand, the aspect ratio (height/radius) of small nascent LDs was always less than 1, i.e., simulated LDs were rather “flat”, indicating that they are not prone to budding. We also found that simulations feature a relatively high surface tension in the bilayer region, not only in the monolayer region, which is compatible with previous experimental results. Overall, our results indicate that simulations of nascent LD systems may be used to provide a microscopic view into the properties of droplet embedded vesicles.

## Methods

### System setup

To generate lipid droplets in bilayer membranes we used molecular dynamics simulations at the coarse-grained level (Martini force field (11) , see next section for details). In the majority of the simulations, we used dioleoylphosphatidylcholine (DOPC) as phospholipid and triolein (TO) as neutral lipid. We initially built phospholipid bilayers containing 2016 or 18144 lipids using the Insane software (12) , then separated the two leaflets and added a layer of TO (325 to 7500 TO molecules) in between them. This resulted in trilayer systems, with a uniform layer of TO sandwiched between two layers of DOPC. The systems were solvated with water particles. The exact composition of each system is reported in Table 1. Hereafter, systems with 2016 phospholipids will be referred to as “small”, the other as “large” systems. After equilibration, these systems rapidly form lens-shaped nascent LDs (Figure 1a). Additional small systems were generated containing 625 TO molecules and different phospholipid types: DPPC, and POPC. Finally, additional small systems were generated using two different neutral lipids, namely trilauryl glycerol (12 carbon atoms) and trioctanoyl (8 carbon atoms) glycerol. Table 1 contains a full list of the simulated systems.

**Table 1.**
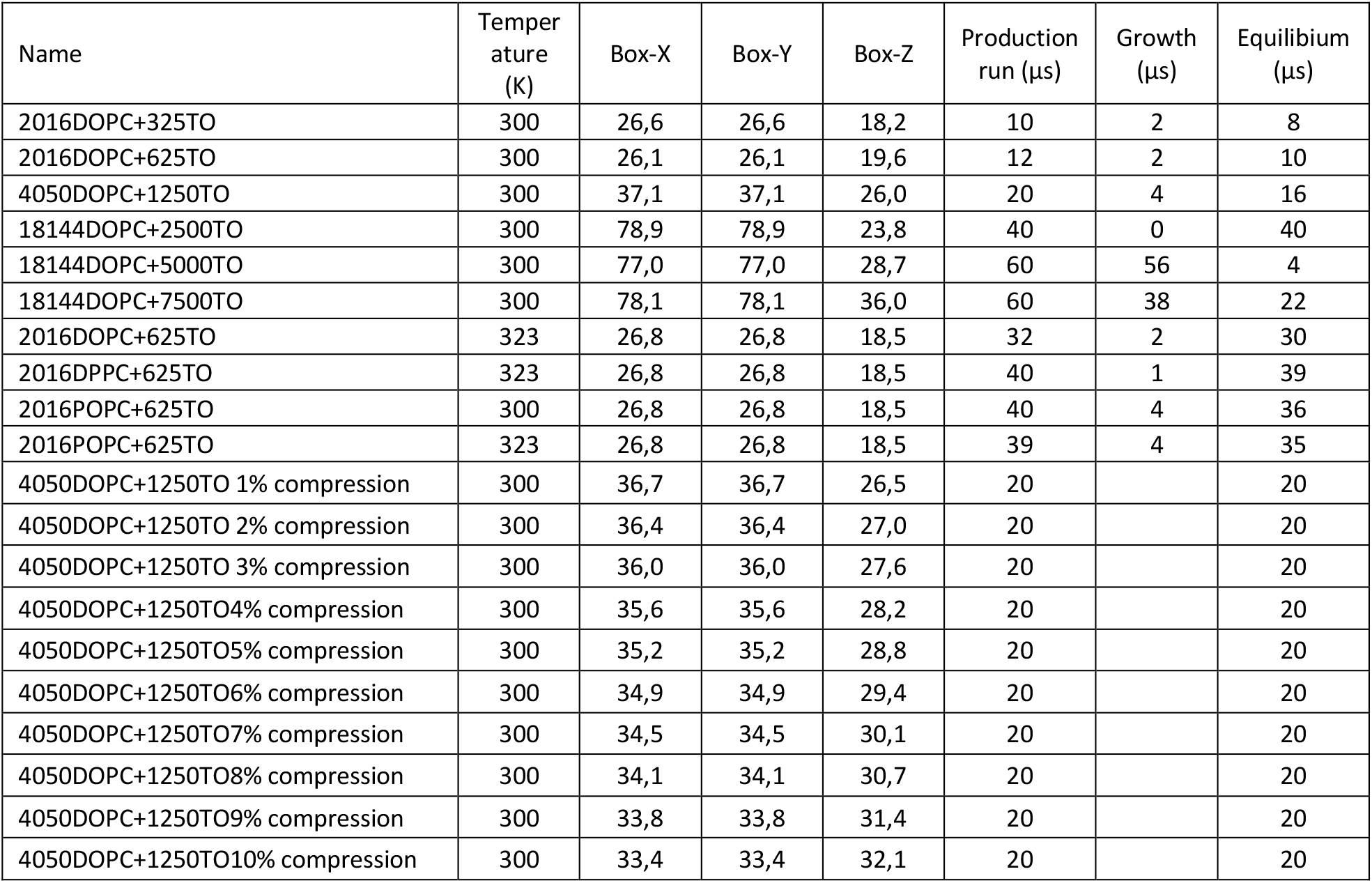
Composition and size of the simulated systems. The growth phase corresponds to 5 τ (where τ is the autocorrelation time of LD size, see Methods), the equilibrium is used for the analysis of shape and equilibrium properties and corresponds to the difference between the total duration of the run and growth phase.

**Figure 1.**
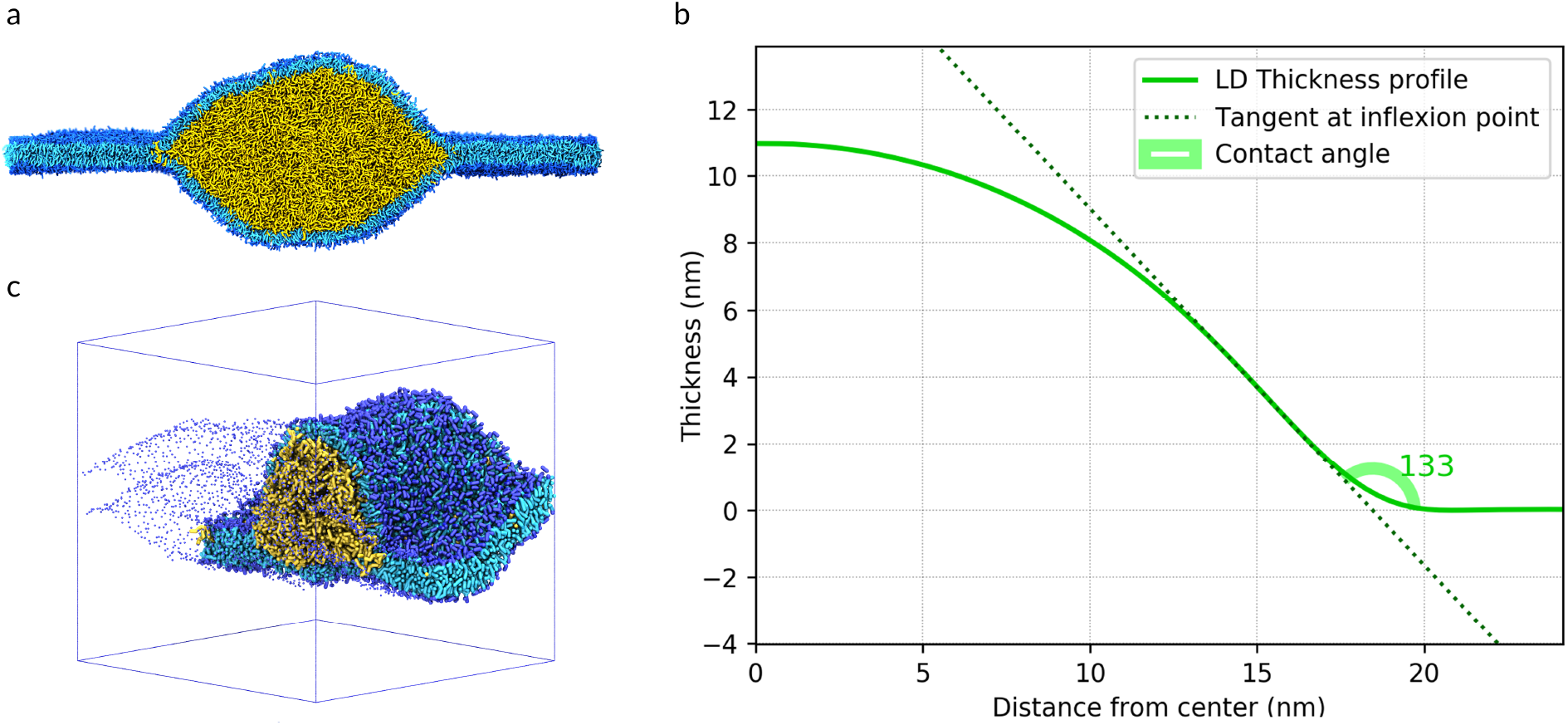
(a) Nascent LD as simulated in this work, and used for the calculation of LD shape and contact angle. (b) Illustration of the methodology to calculate the LD contact angle; the x axis reports the size of the nascent LD; the y axis corresponds to the size of the nascent LD in the direction of the z coordinate (normal to the membrane plane), calculated as specified in the Methods section; for both axes, the value of zero corresponds to the center of the LD. (c) Compressed nascent LD system, cross-section; only half of the LD is shown, to highlight bilayer deformations.

### Simulation details

All simulations were performed with the Gromacs software (v.2016) (13) , using the coarse-grained description provided by the Martini force field (version 2) (11, 14) . The Martini model has been widely used in simulations of various biomolecular systems containing membranes (14) , proteins (14, 15) , nucleic acids (15) , as well as synthetic polymers (16) and nanoparticles (17), and has been shown to provide a realistic description of lipid aggregates and their properties (18).

A cutoff at 1.1 nm was applied for the calculation of non-bonded interactions with the Verlet cutoff scheme, together with the reaction field for electrostatics interactions. The potential was shirted to zero between 0.9 and 1.1 nm with the Potential-shift-Verlet function (13) . The same parameters were used in the calculation of local stresses with the Gromacs-LS software (19).

MD simulations were carried out with the leap-frog integrator and a time step of 20 fs. Temperature was kept constant with the Bussi-Donadio-Parrinello (v-rescale) thermostat (20) (time constant of 1 ps). Pressure was kept constant in equilibration with the Berendsen barostat (21) (time constant of 20 ps, compressibility 4*10^-5^bar^-1^) and in production runs with the Parrinello-Rahman barostat (22) (time constant of 20 ps, compressibility 4*10^-5^bar^-1^). Semi-isotropic pressure coupling was applied in all cases.

Systems were energy minimized using the steepest descent algorithm for 500 steps. After energy minimization, the systems were equilibrated for 10 ns in the NPT ensemble. During equilibration neutral lipids started to phase-separate and formed lens-like nascent lipid droplets embedded in the phospholipid bilayer. Then, production runs were carried out for 10-60 μs for each system (duration reported for each system in Table 1).

Simulations with compression of the nascent LDs were carried out starting from the equilibrium simulation of the 4050 DOPC + 1250 TO nascent LD system. Since this system was run with pressure coupling enabled, we chose our frame by selecting that which had an average x- and y-dimension over the last microsecond of the equilibrium simulation. Using the gromacs .mdp “deform” option, a compression simulation of length 4 µs was then run, compressing the box inwards at a rate of 1nm/µs, with pressure coupling in x and y turned off. As the system began with dimensions 37.195 nm in both x- and y-, this rate of compression resulted in the bilayer being compressed inwards by more than 10% in both dimensions. Frames were then extracted from this simulation at each whole percentage of compression, yielding 10 frames ranging from 1-10% of bilayer compression, from which new equilibrium simulations were started under the same conditions as the original system, only with zero compressibility in the x-y plane to maintain the same surface area.

### Simulation analysis

The number of TO molecules in the LD was monitored over time using our g_aggregate code (23). A modification of this code also allowed us to rewrite the trajectory with the center of mass of the LD translated at the center of the simulation box, resulting in trajectories with a virtually immobile LD. Shape analysis was carried out with our g_thickness code (24) the thickness of the membrane, defined as the distance between the phospholipid heads (PO4 beads) of each leaflet, was computed over an XY-grid, and averaged over time, providing a thickness landscape. We obtained the LD thickness profile by averaging the landscape around its vertical symmetry axis. A python code was developed to extract the geometric properties of the LD shape from the thickness profile. Briefly, the LD shape was fit to a third-degree polynomial function. The radius at which the second derivative of the polynomial approximation cancels out gives the coordinates of the inflection point of the shape curve. The tangent line to the curve at this inflection point was computed, and the intersection of the tangent with the line defining the bilayer was used to estimate the LD radius. The angle between these two lines was defined as “contact angle”, in analogy with the contact angle defined in the theory of wetting/dewetting (see Figure 1b).

The calculation of the local stress tensor from the simulations was performed using the Irving-Kirkwood-Noll procedure, as implemented in GROMACS-LS (19). For multi-body potentials (angles and dihedrals), GROMACS-LS uses the central force decomposition, which provides a unique and symmetric stress tensor for potentials with up to four-body interactions (19). For the local stress tensor grid, we used a cubic voxel of 0.5 nm corresponding to the size of Martini beads. Since our nascent LD systems are large (almost 80 nm lateral size), undulations of the bilayer and monolayer are very substantial; as a consequence, averaging the local stress tensor over the XY plane and over time requires particular care to avoid that undulations cancel out contributions from subsequent time frames and from regions in the XY plane that have different position with respect to the membrane (e.g., different distance from a reference point, such as the center of the bilayer, or the head group region). To this end, we developed a specific code to take into account bilayer and monolayer fluctuations, and extract the pressure profile from the local stress tensor. Averaging of the local stress tensor was carried out using in-house software.

## Results

### Assessment of equilibrium thermodynamics and kinetics

We first generated trilayer systems, with a layer of TO sandwiched between two phospholipid layers (see Methods). Since the TO layer in the trilayer systems were relatively thin, the trilayers were unstable and evolved rapidly by phase separation of TO into a compact droplet, that is the nascent lipid droplet, embedded into the lipid bilayer; part of the TO remained dispersed in the bilayer, indicating that phase separation is dynamic, as expected. Similar observations have been reported before, also based on MD simulations (25–27). Starting from a uniform distribution of TO molecules, we generally observed nucleation of one or more aggregates, and then LD growth during several microseconds. To analyze LD shape, we need to consider only the fraction of simulations representing an equilibrium ensemble, hence we need a quantitative way to assess the equilibration of the system. To this aim, we calculated the size of the largest TO cluster (that is the lens-shaped nascent LD) over time. We observed that, after an initial period of growth, the number of TO molecules in the nascent LDs remained approximately constant, fluctuating around an average value that depends on system composition, size, and temperature. We defined the initial period as the growth phase and the subsequent period as the equilibrium phase. Examples illustrating growth and equilibrium phases are shown in Figure 2.

**Figure 2.**
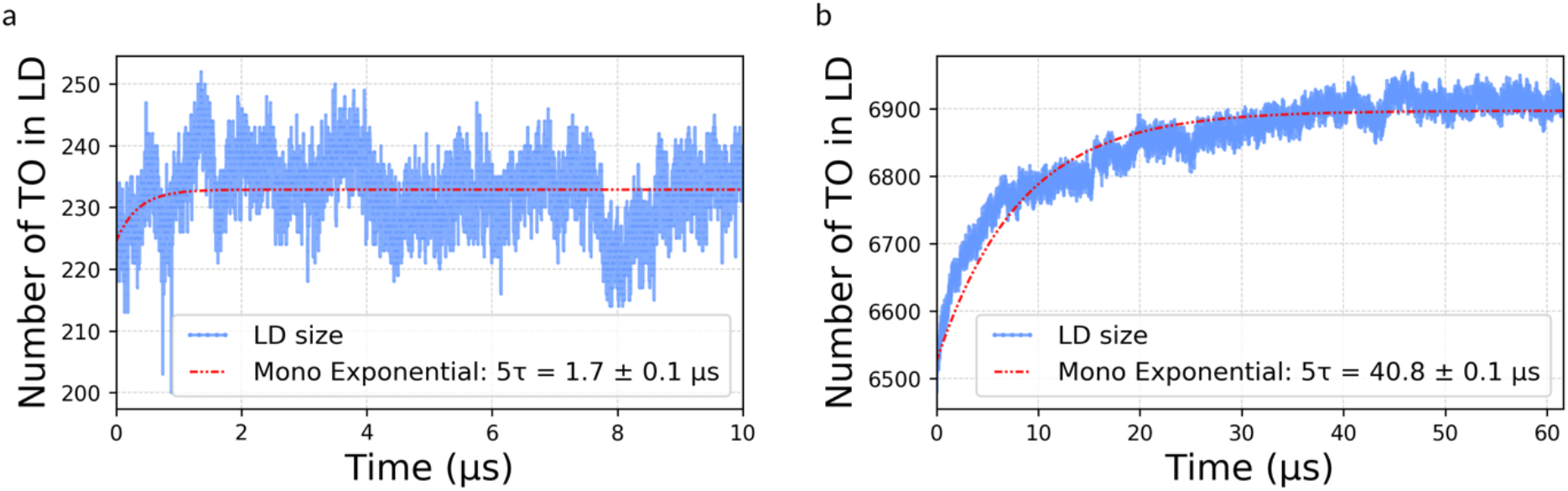
Assessment of system equilibration. (a) Number of TO molecules in the nascent LD in simulation with 2016 DOPC and 325 TO. (b) Number of TO molecules in the nascent LD in simulation with 18144 DOPC and 7500 TO. All analyses were carried out with our g_aggregate code (23).

The duration of the growth phase was different in each simulation, and appeared to depend mainly on the specific initial conditions of the system and the size of the bilayer. As expected, larger systems required much longer equilibration, due to the longer time required for diffusion of TO across the system. For instance, systems of ∼80 nm in lateral size required equilibration times around 50 *μ*s. Our estimate cannot be precise as we simulated only 1 replica of the larger system, due to the large computational cost; therefore, we refrain from providing any quantitative interpretation of growth kinetics. However, available data suggests that the duration of the growth phase does not depend strongly on the number of TO molecules, nor on the nature of the phospholipids (which all form fluid bilayers in the simulation conditions), but rather on the surface area of the system, suggesting that equilibration times are mostly limited by diffusion. Having estimated the equilibration time for each simulation of interest, we extended the duration of the simulation beyond the growth phase (see Table 1). We then analyzed the shape of the nascent LD only considering the equilibrium phase of the simulations, when the number of TO molecules in the nascent LD is approximately constant.

### Simulations are in qualitative agreement with the theory

One of the effects predicted by theory is that LDs with an increasing volume should become more spherical (4). One parameter that indicates whether the LD shape is close to the spherical shape is the curvature of the LD profile: constant curvature is characteristic of spherical shapes. Another parameter indicating similarity to a spherical shape is the LD aspect ratio, i.e., LD height over LD radius (4). This is also related to the contact angle, in analogy with dewetting theory (8). Here we define the contact angle as the angle formed by the membrane plane with the tangent to the droplet profile at its inflection point (Figure 1b). The contact angle decreases (to approach the value of 90 degrees, in the case of a symmetric bilayer) as the LD aspect ratio increases.

We calculated LD curvature, aspect ratio, and contact angle for six nascent LDs with increasing volume and the same phospholipid composition, namely pure DOPC. Shape analysis showed that the curvature is constant at the center of the spherical caps for all simulated LDs; for large LDs, the curvature was constant over a large fraction of the LD surface (Figure 3c-e). Constant curvature indicates that the LD shape is controlled by surface tension of the monolayer region (4), that become prevalent already at relatively small sizes (20 nm in diameter). At the same time, the aspect ratio of all simulated nascent LDs was small (<1) and the contact angle was large (>90 degrees), indicating that the simulated droplets are not close to budding. The contact angle decreased with increasing LDs size, confirming that larger LDs are closer to the budding transition, as expected (Figure 3b). Such change in shape is due to the increase in surface energy relative to elastic energy, as the radius of the LD increases. At large enough sizes, the contribution of bending energy to the shape of the LD becomes negligible compared to the contribution from surface tension, and we expect that the main contribution to LD shape comes from the increased surface energy. These findings agree well with theoretical predictions. We also noticed that, beyond the spherical cap region, LDs are “stretched” or “flattened”, suggesting the existence of forces pulling on the edges of the nascent LD. We will explore such forces later in the manuscript.

**Figure 3.**
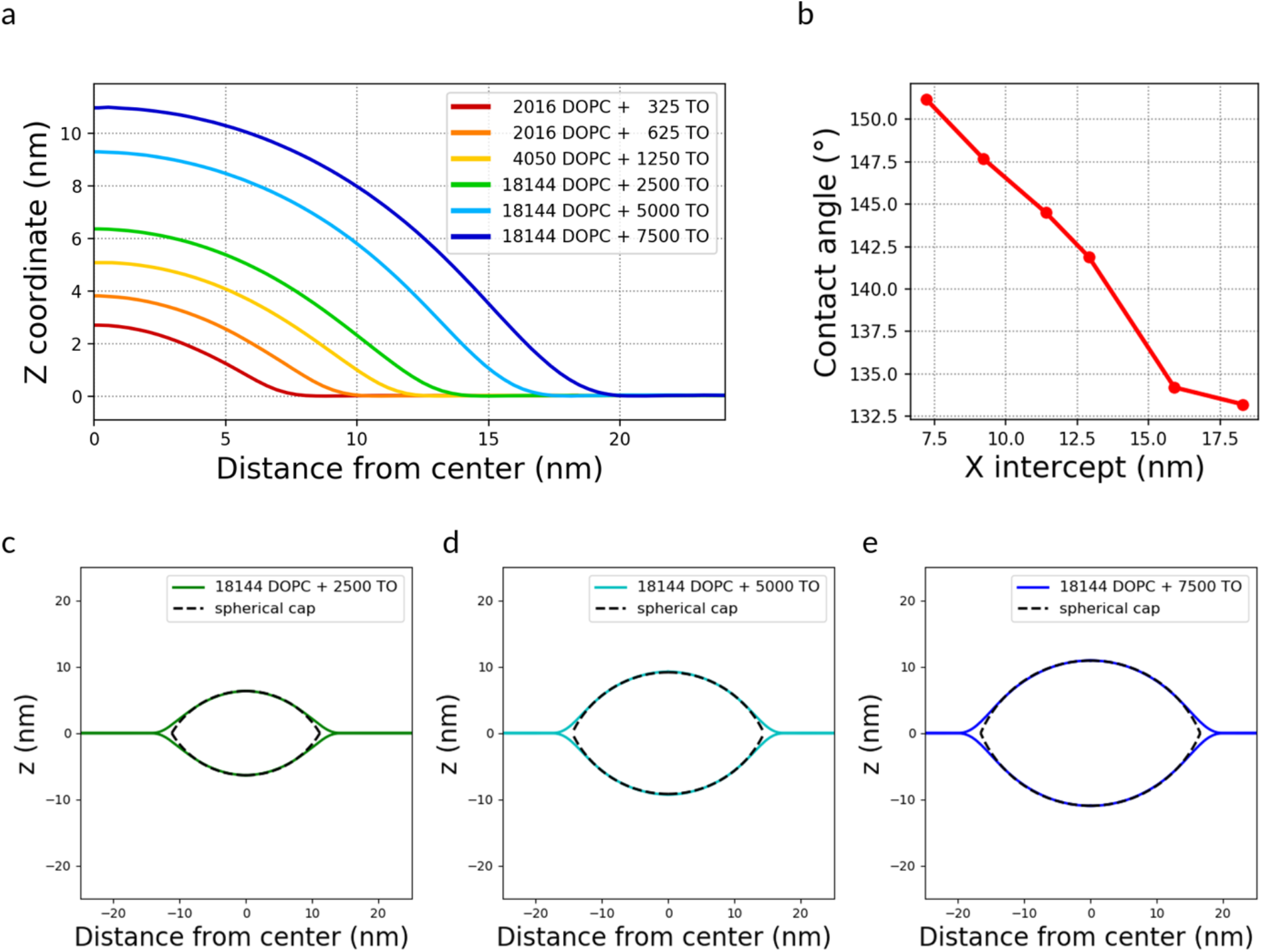
Shape analysis for nascent LDs. (a) Shape profile, after averaging around the axis of symmetry of the LD and over time, for LDs of different size. (b) Contact angle for the nascent LDs in panel B, as a function of the size (expressed as the x intercept).

LD shape theoretically depends on the interplay between the monolayer bending modulus and surface tension (4): higher monolayer surface tension favors spherical shapes, while higher rigidity disfavors them. We therefore aimed to quantitatively evaluate the contribution of each of these factors to LD shape. Measuring the bending moduli of phospholipid monolayers at the TO/water interface is problematic, as the values should depend on the level of coverage of the monolayers (related to their surface tension) and on the amount of TO dissolved in the monolayer. Calculations of the same quantities from MD simulations are also problematic: computational methods to calculate the bending modulus are well tested for lipid bilayers (28), not on monolayers. However, it is reasonable to assume that, to a first approximation, the trends in bending moduli vs lipid composition should be the same for bilayers and monolayers (i.e., if a bilayer consisting of lipid X has bending modulus larger than a bilayer made from lipid Y, then the same will be true also for monolayers). We can also assume that the bending moduli of lipid monolayers at the oil-water interface are approximately half of the bending modulus of bilayers in contiguity, with the same chemical composition.

To understand the effect of bending rigidity on LD shape, we simulated nascent LDs with the same size (i.e., same number of phospholipids and TO molecules) and different phospholipid composition, namely DOPC, DPPC, and POPC. Shape analysis (Figure 4a) showed that LDs made from lipid bilayers with a high bending modulus have a higher contact angle (Figure 4b), i.e., LDs embedded in more rigid bilayers have a lower aspect ratio, as predicted by continuum theory (4). Our results suggest that nascent LD are closer to a spherical shape when formed in softer membranes.

**Figure 4.**
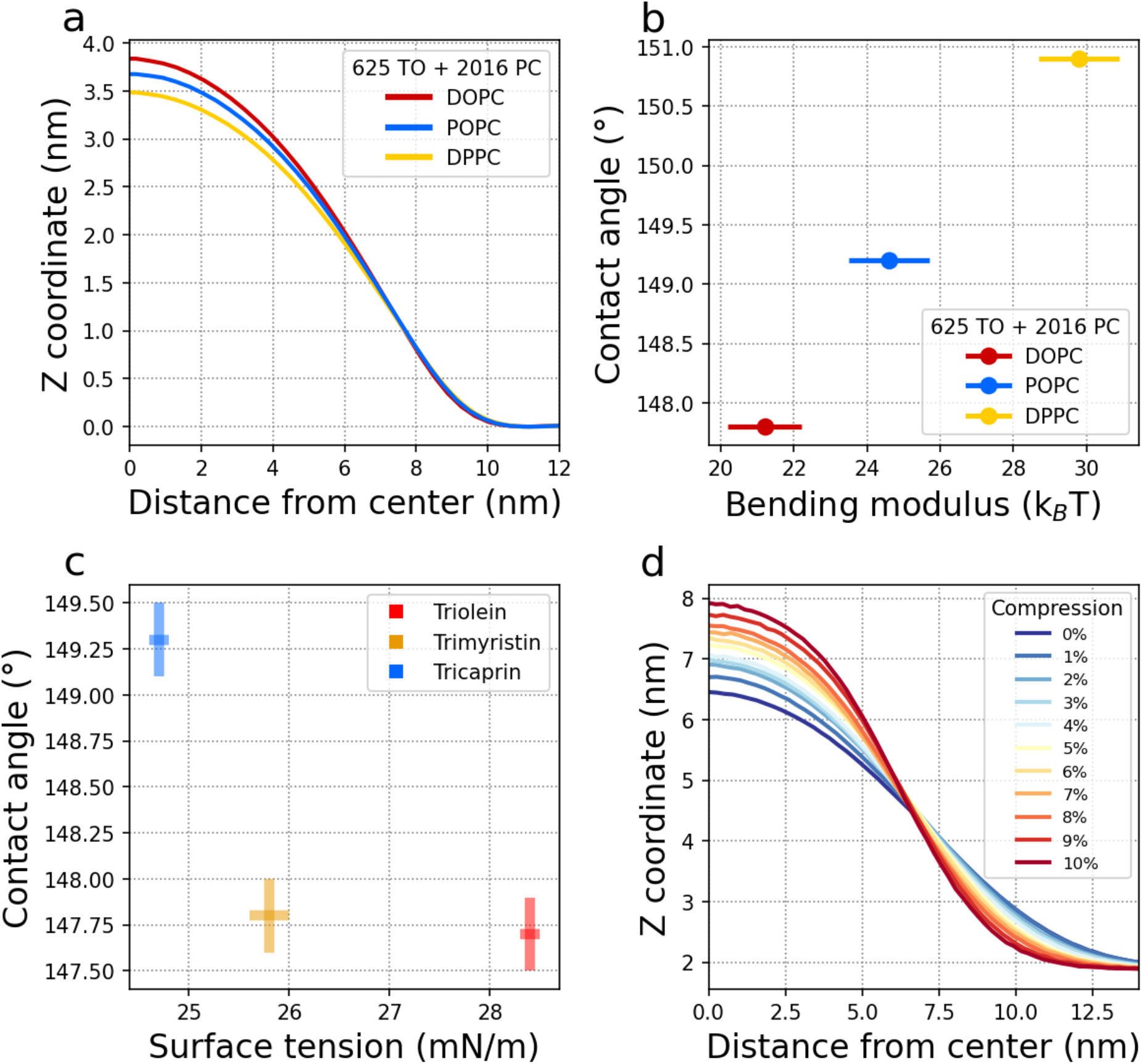
LD shape and bilayer rigidity. (a) LD shape profiles for LDs with the same number of neutral lipids and phospholipids, but different nature of the phospholipids. (b) Contact angle as a function of pure lipid bilayer bending modulus of the bilayer, as calculated in ref. (28). The DPPC systems were ran at 323K, while all others at 300K. (c) Contact angle as a function of surface tension of the oil-water interface. (d) Shape analysis of nascent LD systems, with the bilayer compressed in the membrane (XY) plane; the degree of compression is indicated as the ratio of the actual X dimension vs the X dimension in the uncompressed simulation.

To understand the influence of surface tension on LD budding, we simulated nascent LDs containing different triglycerides, having different interfacial tensions with water. We chose trimyristin and tricaprin, which have a lower tension than triolein. The simulated nascent LDs contained the same number of neutral lipids and phospholipids and the same phospholipid composition. Shape analysis showed a correlation between surface tension and contact angle: nascent LDs with lower surface tension have lower aspect ratio and higher contact angle (Figure 4c). This is a small effect in terms of contact angle, and it is difficult to comment on its biological relevance. However, we note that TO, the most abundant neutral lipids present in LDs in cells, has a higher surface tension with water, and LDs composed of TO are slightly more prone to budding.

We also investigated the effect of compressing the bilayer in the XY plane (Figure 4d). Lateral compression reduces the surface tension of the bilayer by increasing the surface density of the lipid bilayer. Equilibrium simulations of compressed nascent LDs show a clear and consistent increase in the LD aspect ratio, as expected, leading to more spherical droplets.

### Quantitative comparisons between simulations and continuum theory

The shape profiles of nascent LDs simulated with our coarse-grained model comply qualitatively with the predictions of continuum theory, but do they reproduce quantitatively those predictions? We addressed this question by fitting the shapes calculated from our simulations with theoretical shapes from ref. (4). To this end, we used the same assumptions as in the theoretical analysis; in particular, long-range molecular interactions contribution to the spontaneous curvature of the monolayers were neglected, simplifying the theoretical shape equations. Under those assumptions, the LD shape equation (equation 1) contain only 3 parameters: the elastic length *λ,* defined as *λ* =(*γ*_m_/*γ*_m_)^1/2^, V/*λ*^3^ (where V is the LD volume), and *γ*_b_/*γ*_m_ (surface tension of the bilayer and the monolayer). The volume of the droplet being known, we used *λ* and *γ*_b_/*γ*_m_ as fitting parameters. Values of surface tension for the monolayer (*γ*_m_) can be estimated from *λ* (that is obtained from the fitting procedure), with some assumptions on the bending rigidity of the monolayer (*γ*_m_). Bending rigidities have been measured at ∼20 k_B_T for pure DOPC bilayers in numerous studies, using different experimental and simulation methods (28–30). Furthermore, Santinho *et al*. showed that the addition of a small amount of TO decreased the bending modulus by nearly a factor of 2, from 21 +/-4 k_B_T to 12 +/-4 k_B_T (31). Assuming that the bending rigidity of monolayers is close to half of the value for the bilayer (32) , we estimate an approximate value of *γ*_m_ = 6 k_B_T. This is in good agreement with values estimated for DOPC monolayers (32).

As described in the Methods section, simulations were carried out using the semi-isotropic pressure coupling method, with pressure of 1 bar in the XY plane and in the Z dimension. The same setup is generally used in simulations of bilayer membranes, where it provides zero surface tension for the membrane system. Based on the expectation of a tensionless bilayer, we first fitted the simulation-based shapes imposing zero surface tension in the bilayer, i.e., using a single fitting parameter, *λ.* In this case, the results were unsatisfactory: the simulated shapes had approximately constant curvature in the central region, while theoretical shapes did not (Figure 5a). Indeed, in the absence of bilayer tension, shapes with constant curvature over a large fraction of the LD surface (obtained from the theory) would have an aspect ratio close to one. Several possibilities exist to explain the mismatch between simulated shapes and theoretical shapes: (a) the surface tension is actually non-zero in the simulations, both in the monolayer and in the bilayer regions; (b) the theory is inadequate to describe our nano-sized system. We do not comment here on the possibility that the theory breaks down for our specific systems, and in the following we explore the other possibility.

**Figure 5.**
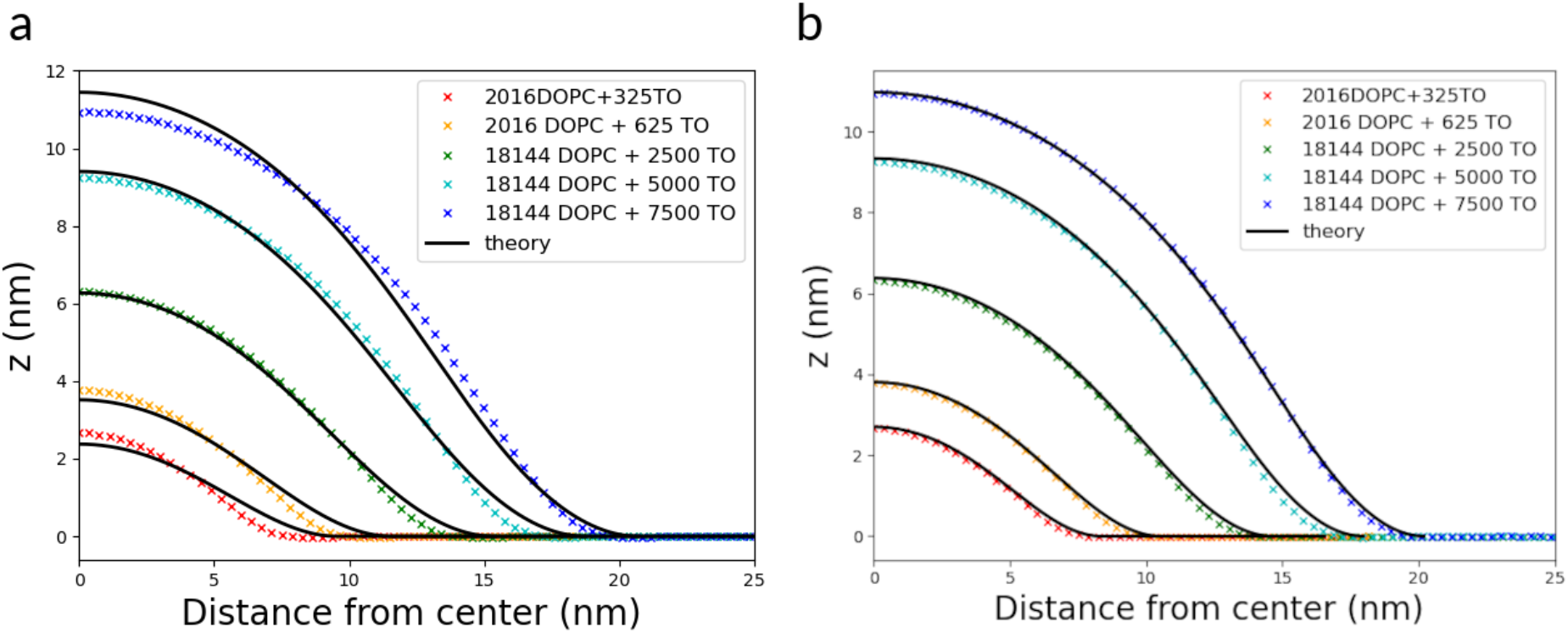
Fitting the simulated shape profiles with the theoretical shape equation. Continuous lines represent the shape predicted by the theory, symbols the simulated shape. (a) Fit obtained by imposing zero surface tension in the bilayer. The fitting procedure yields *λ* = 6.877 nm, from which we calculate *γ*_m_ = 0.51 mN/m. (b) Fit obtained with *λ* = 3.308 nm, and *γ*_b_*/γ*_m_ = 0.834 (7500 TO), 0.844 (5000 TO), 0.944 (2500 TO), 0.956 (625 TO), 0.824 (325 TO).

Given that the semi-isotropic pressure coupling scheme, which assumes homogeneity in the XY plane, is not strictly valid in the presence of the oil droplet, we first considered the possibility that the bilayer tension is not vanishing in our system, and fitted the simulated shapes using both *λ* and *γ*_b_/*γ*_m_ as variables (i.e., two fitting parameters). In this case, the fit was much more accurate (Figure 5b). Assuming the same value of bending rigidity as above (*κ*_m_ = 6 k_B_T), we obtained *γ*_m_ = 2.19 mN/m and then *γ*_b_ close to 2 mN/m for all LD volumes. The value of the bilayer tension obtained from the fit seems surprisingly large, considering that in the absence of nascent LD the bilayer would be strictly tensionless, and knowing that the tension in the ER membrane (where LDs are generated) is very small, nearly vanishing (33). However, this is not surprising if we consider that the monolayer tension is also higher than experimentally measured in micrometer-sized droplet embedded vesicles: in the case of droplet embedded vesicles made of DOPC and TO, the monolayer tension *γ*_m_ is about 1.4 mN/m when the bilayer is tensionless (34). In experiments on droplet embedded vesicles, bilayer and monolayer surface tension are linearly related (34) (*γ*_m_ = 0.45*γ*_b_ + 1.32 mN/m), and a monolayer tension of 2.19 mN/m would correspond to a bilayer tension of 1.93 mN/m, very close to the one predicted by fitting the simulated LD shapes. Hence, the present findings are fully compatible with available experimental data on droplet embedded vesicles with non-vanishing surface tension (34) and with the theoretical model (4).

### Surface tension from the lateral pressure profile

Fitting the curve from simulations with the theoretical curve indicates that significant surface tension should be present in the simulated systems, both in the monolayer and in the bilayer region. To confirm this, we calculated directly the mechanical tension by analyzing the local stress tensor. In materials undergoing strain, neighboring particles exert forces on each other to counteract deformations. The stress tensor σ describes these internal resistive forces. In simulations, we can calculate the local stress tensor directly from kinetic energy and the virial, i.e., from velocities, positions, and forces in the system (35). From the local stress tensor, in simulations of flat, periodic lipid bilayer systems, we can then calculate the lateral stress profile σ_L_, describing the internal forces in the bilayer as a function of the position along the membrane normal; this is defined as

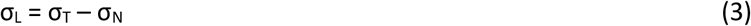

where σ_L_ is the lateral stress, σ_T_ is the tangential stress (i.e., the stress in the plane of the membrane), and σ_N_ is the normal stress (i.e., in the direction normal to the membrane). In the case of a flat membrane oriented with the normal in the direction of the Z axis, this can be written as:

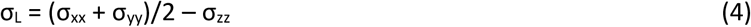

where σ_xx_, σ_yy_, σ_zz_ are the diagonal components of the stress tensor. The stress tensor can also be calculated separately in sub-cells of the simulation cell, expressed as a function of the Z coordinate, and averaged over the simulation time and over the XY plane (Figure 7a). In this case, the lateral pressure profile π(z) is often reported (instead of the stress), calculated as:

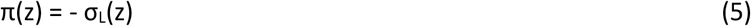

**Figure 7.**
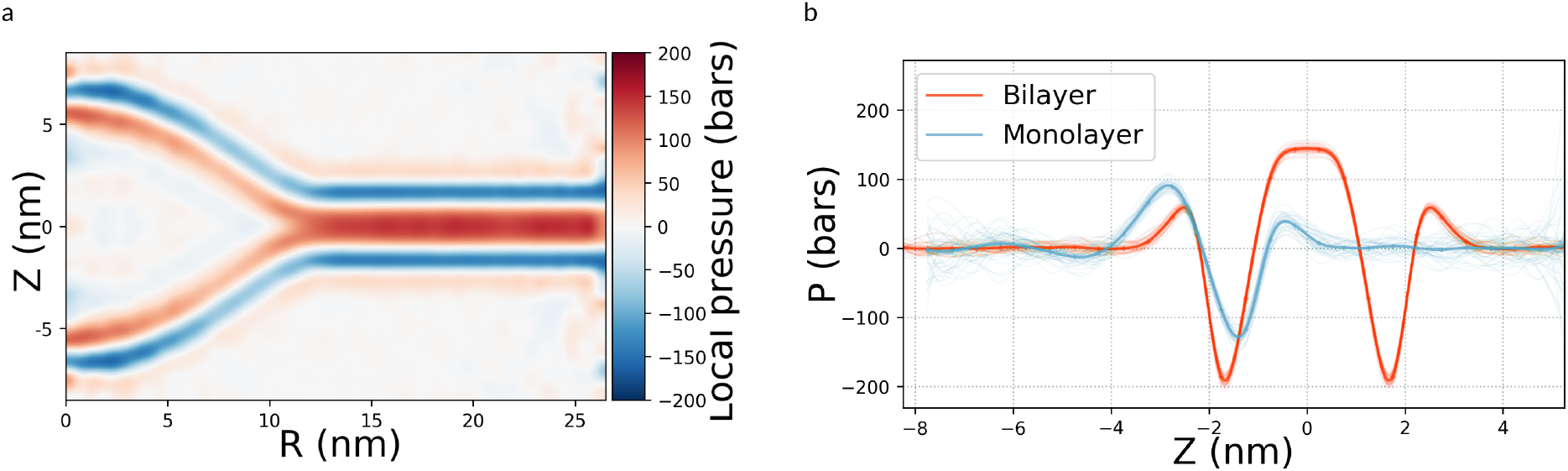
(a) Average lateral pressure in a nascent LD system (4050 DOPC+1250 TO). The characteristic negative peak corresponds to the head groups and positive peak to the acyl chains. (b) Pressure profile in the monolayer (blue) and the bilayer (red) region as a function of the distance from the center of the bilayer (z=0) or the monolayer-oil interface. For the bilayer, the pressure profile (red line) is plotted with z=0 at the bilayer midplane, while the profile for the monolayer region (blue) is plotted with z=0 at the average position of the terminal hydrophobic beads of the lipid tails.

The lateral pressure profile has been reported in the literature for a number of different bilayer systems (19, 36, 37), and it typically presents a characteristic large negative peak corresponding to the interface between the hydrophobic interior of the membrane and the hydrated head group region (close to the glycerol backbone, in phospholipids), and smaller positive peaks in the head group region and the membrane interior (19, 36, 37). In the case of a flat membrane, the surface tension matches exactly the mechanical tension obtained from the simulations; if the membrane normal is oriented as the Z axis, the surface tension can be computed easily by integrating the pressure profile over the membrane thickness:

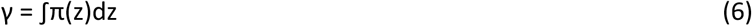

Similarly, the surface tension can also be calculated easily from the local stress tensor in systems with spherical symmetry, such as (spherical) liposomes, or even cylindrical symmetry (19, 36, 37). However, the nascent LD systems simulated here do not have planar, nor spherical, nor cylindrical symmetry, and calculations of local stress or lateral pressure turned out to be problematic, because the large undulations in the systems (which have lateral size of ∼80 nm) cause large uncertainties, with sampling up to 20 *μ*s. Calculation of the local stress tensor in each given time frame is straightforward, but interpretation in terms of surface tension requires averaging over the region corresponding to the bilayer plane (that is parallel to the XY plane, and has planar symmetry) and over the monolayer region corresponding to the spherical cap (see Figure 7); averaging causes significant “blurring” of lateral pressure due to undulations, and such effect increases with the size of the system (just like undulations increase with system size). Our best estimates indicate that the surface tension in the bilayer region is indeed positive and large (Table S2), of the order of a several mN/m, beyond the statistical uncertainty, in good agreement with the predictions obtained by our fitting procedure (Figure 5).

## Discussion

The shape of nascent lipid droplets can be considered as a proxy for their tendency to bud out of the membrane. According to an established theory (4), the shape of nascent LDs should depend on the LD size and the chemical nature of the oil and the phospholipids. Here we simulated nascent LDs, i.e., isolated TO droplets embedded in lipid bilayers, with different size and different chemical composition, calculated parameters related to their shape, and compared the shape of the simulated LDs with the shapes predicted by the theory. We found good agreement between simulations and theory, with LDs becoming more spherical (a) as they grow in size, (b) as their oil-water surface increases, (c) as the phospholipids form softer membranes, and (d) the bilayer is compressed, reducing surface tension. Also, we noticed that the LD shape resembles a spherical cap, with constant curvature over a large fraction of the LD surface, already for relatively small nascent LD sizes (10 nm in radius). This indicates that the shape is mostly determined by surface tension, more than by membrane bending rigidity, as predicted by theoretical work (4)(5). We note, however, that constant curvature alone does not unequivocally prove tension-dominated behavior, as spherical shapes can arise from bending energy alone under certain boundary conditions. The more definitive evidence comes from the elastic length 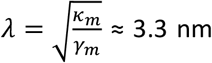 extracted from our fits, which is significantly smaller than the radii of the droplets studied here (5–20 nm). This places the system firmly in the tension-dominated regime, where surface tension is the primary determinant of the droplet shape.

We then fitted the simulated LD shapes in order to interpret simulation results in terms of elastic properties of the systems. Excellent quality fits were obtained when using monolayer and bilayer surface tensions as free fitting parameters. In this case, the fitting procedure predicted monolayer and bilayer surface tensions of the same order of magnitude, both around 2 mN/m, with the monolayer tension only marginally higher than the bilayer tension. We note that similar values of surface tension in the monolayer and bilayer region are predicted also by the empirical relationship for analogous systems in ref. (34) (see Figure 2C in that reference), for a monolayer tension of 2 mN/m. The existence of high surface tension in the bilayer region was also confirmed by calculations of the local stress tensor, which are completely independent of the fitting procedure. Despite significant statistical uncertainties, our results consistently indicate that the shape of simulated LDs results from non-vanishing surface tension in the bilayer region.

A third, independent argument substantiating the non-vanishing bilayer tension emerges from the scale-dependence of the contact angle. Although the apparent contact angle measured at the inflection point decreases systematically with increasing LD volume (Fig. 3B), the mechanical tension ratio γ_b_/γ_m_ extracted from the full-shape fitting remains constant across all simulated sizes. This apparent dichotomy is rationalized by the theoretical framework of Lipowsky et al. (38), which rigorously demonstrates that while apparent (geometrically-defined) contact angles vary with vesicle size and global geometry, the intrinsic contact angle—and hence the underlying surface tension ratio—is a genuine material property, independent of system dimensions (38). The invariance of our fitted γ_b_/γ_m_ thus qualifies the associated bilayer tension as an intrinsic mechanical parameter of the simulated interface, rather than a size-dependent artifact arising from finite-size effects or curvature coupling. This third line of evidence, fully independent of both the quality of the shape-fitting procedure and the local stress calculations, reinforces our conclusion that the bilayer surrounding the nascent LD sustains a finite, non-zero surface tension.

Large values of the surface tension (∼2 mN/m) in bilayer membranes may appear as surprising, since we impose zero tension on the overall simulation system (via the pressure coupling algorithm). However, in simulated systems, pressure coupling imposes zero tension on the overall simulation box but not on individual monolayer and bilayer regions. Considering periodic systems connected at the boundaries, a valid alternative for simulations of nascent LD is the use of fixed area in the XY plane, as done by Bahrami and co-workers (39). Here we did carry out additional simulations at fixed area, using the same area obtained from simulations with semi-isotropic pressure coupling; we obtained exactly the same LD shape, hence the same tension in bilayer and monolayer, as expected. It is also possible to laterally compress (or expand) the system and obtain higher (or lower) surface densities of phospholipids, and simulate them at fixed area; simulations with different values of the area show that LD shapes depend essentially on the surface density of the phospholipids (see Fig. 4d), as already noticed in ref. (39). We notice that varying the area in simulations is analogous to varying bilayer tension in droplet embedded vesicles, e.g., by micropipette aspiration; in both cases, different LD shapes are produced (40, 41).

While large values of surface tension in the bilayer can be reproduced experimentally in model systems (e.g., droplet-embedded vesicles), they do not correspond to the situation in the ER, where bilayer tension is two orders of magnitude lower (∼0.01 mN/m) (33). This is an important conclusion, as it indicates that the standard setup used in simulations of nascent LDs (42–49), with flat periodic systems and semi-isotropic pressure coupling, is not suitable for reproducing the surface tension (and related properties) of nascent lipid droplets in the ER, or in any experimental system with negligible or low surface tension in the bilayer. Simulations at fixed area (39) are a valid alternative, but demand an iterative procedure, calculating the surface tension for each area until the desired (experimental) surface tension is reproduced. Also, such approach would predict the shape of LDs given a certain tension in the bilayer, but would not be predictive of the tension in ER tubules. A second alternative is to simulate systems with non-periodic membrane boundaries, for instance vesicles, as shown by Voth and coworkers (50, 51); in this case, membrane tension would be imposed by the volume of water enclosed within the vesicle or tube, because water permeation through the membrane is slow on the simulation time scale. Such constraint can be relaxed if the vesicle has pores, yielding identical pressures on both sides of the membrane; in this case, we would expect low membrane tension, stemming only from membrane curvature. Despite the difference in morphology, such non-periodic vesicular systems would provide a closer proxy to lipid droplets in the ER, at least in terms of surface tension, but they are more complex to build and computationally more expensive.

## Conclusions

We studied the shape of lipid droplets as obtained from MD simulations at the coarse-grained level, and compared it to the predictions by an established theory (4) . We identified the parameters affecting LD shape: LD volume, as larger LDs are more spherical in shape; membrane softness, as LDs embedded in less rigid bilayers are more spherical; surface tension between oil and water, as a higher surface tension makes LDs more spherical. These trends are consistent with theoretical predictions, indicating that the predictions by continuum theory are valid for nascent LDs even at very small scales. Moreover, LD shapes calculated from simulations can be fitted with the theoretical shape equation in ref. (4), providing an estimate of the surface tension in the monolayer and in the bilayer region. Interestingly, the fit indicates that, in the simulations, the surface tension in the bilayer region is non-vanishing, despite applying the same pressure in the membrane plane and along its normal. This result is also confirmed by local stress calculations, as well as theoretical considerations. We conclude that simulations of lipid droplets embedded in flat, periodic membranes do not reproduce the surface tension of LDs in tensionless ER membranes. However, they do reproduce the properties of droplet embedded vesicles in the presence of bilayer tension. Moreover, simulations carried out at fixed area, with different levels of lateral compression, do produce different LD shapes, with aspect ratios closer to 1 for highly compressed membranes; varying the level of compression in simulations is analogous to varying the tension in experiments with droplet embedded vesicles. Overall, the shape of LDs observed in simulations is in excellent agreement with available theories, and simulations of nascent LD systems may be used to provide a microscopic view into the properties of droplet embedded vesicles.

## Acknowledgement

LM acknowledges Samuli Ollila, Markus Deserno, and Sonia Cambiaso for fruitful discussions. Molecular dynamics calculations were carried out at CINES, GENCI grant no. A0080710138, A0100710138, A0120710138, and A0140710138 to LM. VN, JC, and LM thank CC-IN2P3 (https://cc.in2p3.fr) for computing services (data storage and backup). LM acknowledges funding from the *Institut national de la santé et de la recherche médicale* (INSERM). This work was supported the ANR-NANODROP (ANR-17-CE11-0003) and ANR-LIPRODYN (ANR-21-CE11-0032) to ART and LM.

## Author contributions

LM, LF, and ART designed the study. VN, JC, and LM performed and analyzed all simulations. FD and LF performed the fitting of simulated LD shapes with theoretical shapes. VN, JC, and LM wrote the manuscript, with contributions from all authors. All authors discussed and approved the final version of the manuscript.

