## Supporting Information for "Lipid droplet shape and tendency towards budding: insight from theory and molecular simulations"

Vincent Nieto[1], Jackson Crowley[1], Francois Deslandes[2,3], Abdou Rachid Thiam[2], Lionel Foret[2,*], Luca Monticelli[1,4,*]

[1] Molecular Microbiology and Structural Biochemistry (MMSB), UMR 5086 CNRS, Université Claude Bernard Lyon 1, F-69007, Lyon, France

[2] Laboratoire de Physique de l'École Normale Supérieure, ENS, Université PSL, CNRS, Sorbonne Université, Université de Paris Cité, F-75005 Paris, France

[3]Université Paris-Saclay, INRAE, MaIAGE, 78350, Jouy-en-Josas, France

[4] Institut National de la Sante et de la Recherche Medicale (INSERM), France

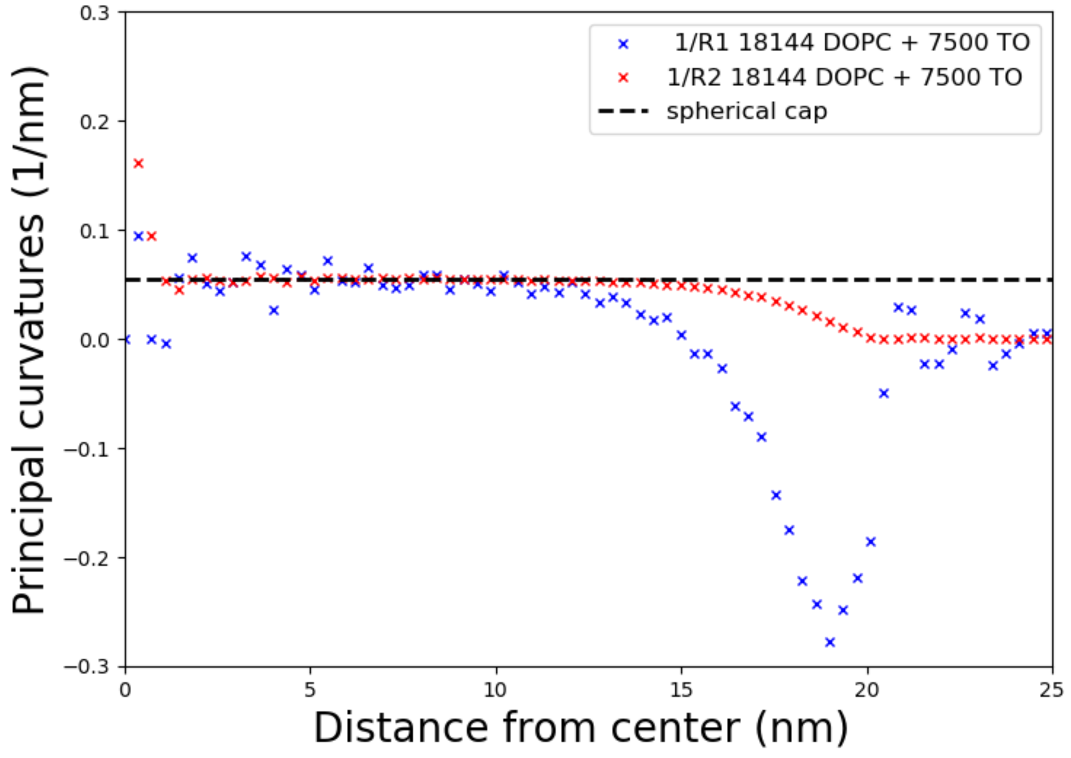


Figure S1. Principal curvatures of simulated nascent LDs containing 7500 TO lipids.

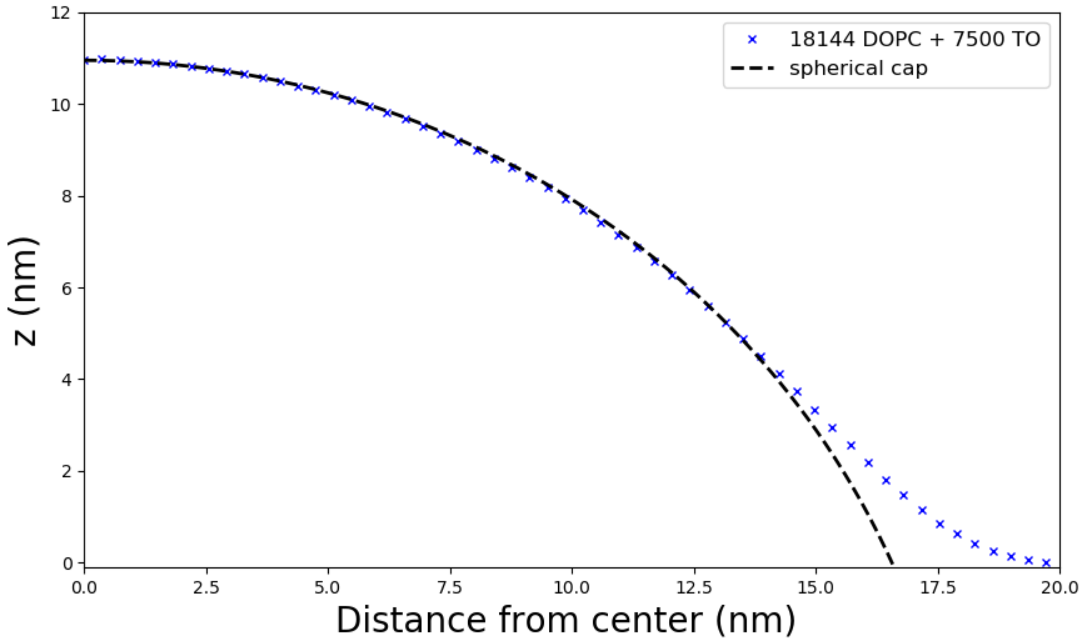


Figure S2. Fitting the simulated LD shape with a spherical cap, for simulated nascent LDs containing 7500 TO lipids.

| Nascent LD – system description | Bilayer surface tension $\gamma_B$ (mN/m) |
|---|---|
| 2016 DOPC+325 TO | 8.2 ± 4.5 |
| 2016 DOPC+625 TO | 10.7 ± 3.5 |
| 4050 DOPC+1250 TO | 6.6 ± 3.6 |
| 18144 DOPC+2500 TO | 12.6 ± 5.5 |
| 18144 DOPC+5000 TO | 4.9 ± 4.3 |
| 18144 DOPC+7500 TO | 6.1 ± 4.7 |

Table S1: Bilayer surface tension computed used the local stress in the simulated LD systems.
